# Liposome Encapsulation Enables Near-Native Cryo-EM Structural Determination by Shielding Macromolecules from Non-Physiological Interfaces

**DOI:** 10.1101/2025.05.06.652550

**Authors:** Zikai Gao, Lijuan Ma, Hang Fu, Chenguang Yang, Dongfei Ma, Hongtao Zhu, Yumei Wang, Dapeng Sun, Shuxin Hu, Chunhua Xu, Wei Ding, Jianbing Ma, Ying Lu, Ming Li

## Abstract

Cryo-electron microscopy (cryo-EM) has emerged as a very powerful tool for high-resolution structure determination of macromolecule complexes. However, sample preparation remains a major bottleneck in cryo-EM workflows. Most structures were resolved not in a native solution environment, but rather in the non-physiological interfaces such as the air-water interface (AWI), where the folding/unfolding energy landscape of macromolecules may be significantly altered, leading to artifact, preferential orientation, particle disassembly or even denaturation. To address this challenge, we developed a robust, sample-independent method utilizing liposome encapsulation that preserve macromolecules in a solution-like and near native environment throughout sample preparation. Using equine spleen apoferritin and the *Escherichia coli* (*E. coli)* ribosome as model systems, we demonstrate efficient particle incorporation into liposomes and successful high-resolution structure determination. Notably, our method yields significantly reduced particle disassembly, denaturation, and preferential orientation compared to samples on holey carbon and graphene grids. We anticipate that this approach will facilitate structural studies of other challenging macromolecular complexes that are sensitive to interfacial effects.

## Introduction

Over the past decade, single-particle cryo-EM has become the leading technique for high-resolution structural determination of macromolecules ^1^. Despite its success, a major challenge persists in sample preparation: approximately 90% of macromolecules are attached at the air-water interface (AWI) during conventional cryo-EM grid freezing (Figure 1a, right panel) ^2^. The AWI imposes an energy landscape distinct from that of bulk solution, potentially distorting macromolecular conformations away from their native states ^3^. Moreover, exposure to the AWI can induce denaturation, disassembly, and/or preferential particle orientation, making structure determination less efficiency or even unfeasible ^4,5^. In addition, cryo-EM sample preparation requires extensive optimization of multiple parameters (e.g., sample concentration, blotting conditions, ice thickness), which are highly sample-dependent and time-consuming.

**Figure 1.**
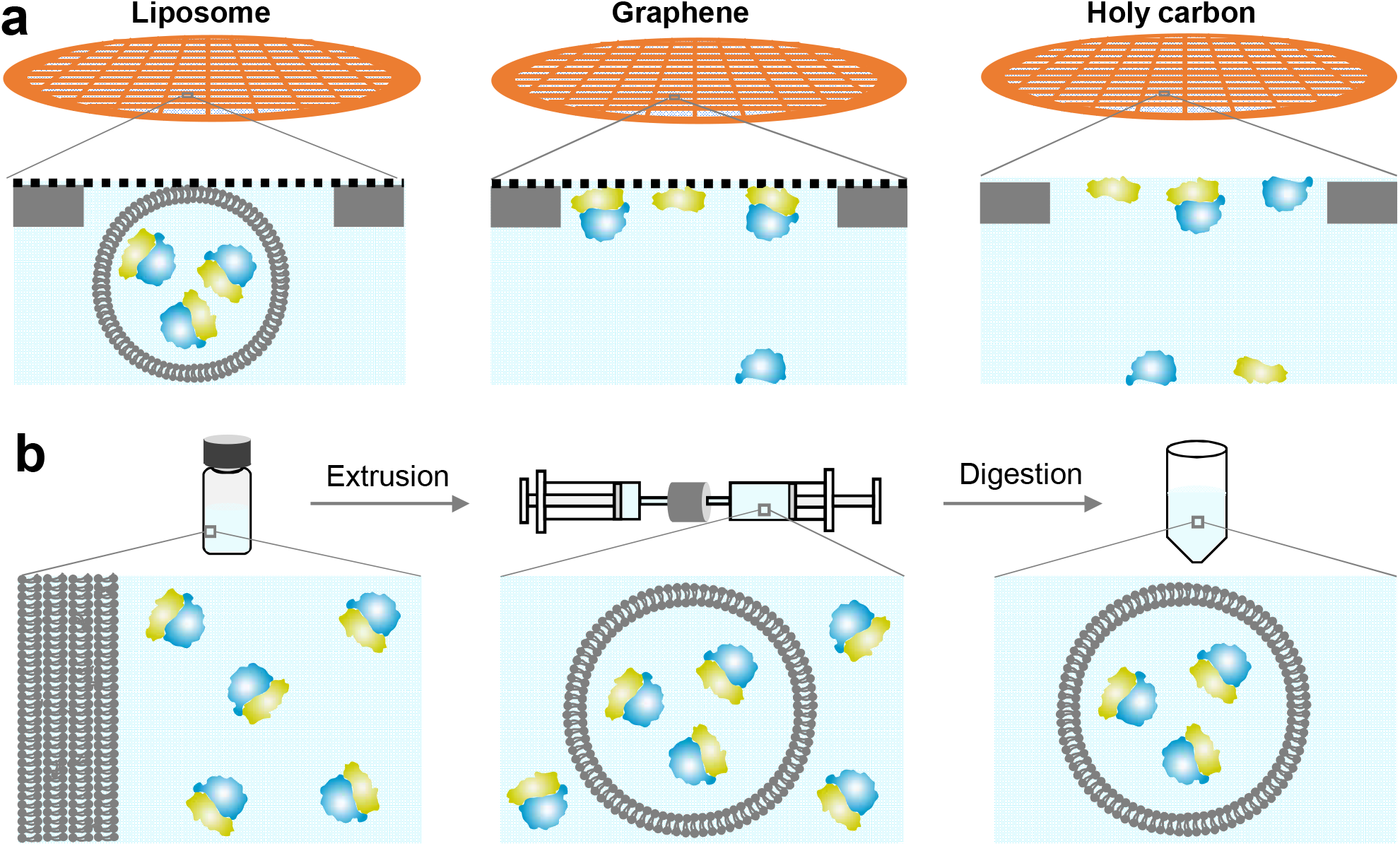
Liposomes encapsulating method in relative to other sample making methods. **a**, Cartoon illustrations of the position of particles in the holes by various sample preparation methods. **b**, A schematic illustration of the sample prepration steps to encapsulate macromolecule in liposomes.

To mitigate these challenges, several strategies have been explored ^6^, including detergent, chemical crosslinking, rapid plunge-freezing ^7^, surface adsorption (Figure 1a, middle panel) ^8-11^, electrospray ^12^, and nanofluidic chips ^3,13^. While these methods have shown promise for several cases, none have proven universally effective or widely adopted. Here, we present a simple, structure friendly, reproducible, and broadly applicable approach: encapsulating macromolecules within protective liposomes (∼100 nm) that adsorb onto graphene-coated grids (Figure 1a, left panel). Macromolecules could be efficiently loaded into liposomes by gentle mixing with lipid suspensions ^14^. To optimize liposome size for cryo-EM, extrusion through defined-pore membranes could be employed (Figure 1b, middle panel). Following extrusion, unencapsulated macromolecules were enzymatically removed, yielding samples ready for cryo-EM imaging (Figure 1b, right panel). Since liposomes preferentially adsorb to carbon rather than suspend in vitreous ice ^15,16^, holy occupancy could be enhanced by depositing a graphene layer over holey carbon grids (Figure S1).

Applying this liposome-graphene hybrid method, we successfully preserved apoferritin and *E. coli* 70S ribosomes from AWI or other structurally disruptive interfaces. Strikingly, nearly all ribosomes remained intact, compared to severe dissociation into 50S and 30S subunits (with only 64% and 28% 70s were kept assembled, respectively) on graphene and holey carbon grids. Furthermore, key ribosomal surface features retained full density in liposome-encapsulated samples, unlike the missing densities observed with conventional grids. Finally, particle orientations were nearly isotropic, in relative to the pronounced preferential orientation seen with standard methods. Collectively, these results underscore the efficacy of our approach in maintaining macromolecular integrity and enabling high-resolution structure determination.

## Results

### Macromolecules could be efficiently encapsulated and randomly dispersed in liposome

To validate the method, two model systems: apoferritin (443 kDa, octahedral symmetry) ^17-20^ and *E. coli* 70S ribosome (2.3 MDa, C1 symmetry) ^21-23^ were selected. These well-characterized complexes represent extremes in size and symmetry. Despite their inherent structural stability, both unincorporated/unprotected complexes were completely digested and removed within hours (Figure S2). Electroneutral lipids have demonstrated excellent passivation properties, effectively preventing nonspecific macromolecular adsorption ^24^. This passivation effect remains robust even in the presence of 2-20% negatively charged lipids ^11,25^. Based on these findings, we formulated lipid compositions with 95% electroneutral lipids as the dominant component.

We systematically evaluated encapsulation efficiency across a concentration gradient (5.4, 14.2, and 50 μM). Particle incorporation showed a positive correlation with sample concentration, with similar incorporation efficiencies observed for both complexes at equivalent concentrations (Figure S3). For high-resolution reconstruction, which typically requires >100,000 particles per day for one single dataset, we determined that a minimum concentration of 10 μM was necessary to achieve ≥10 particles per micrograph. Before cryo-EM data collection, 3D tomography analysis was processed. These two samples revealed distinct particle distributions between our liposome-encapsulated samples and conventional preparations (Figure 2, S4). For the liposome-encapsulating samples, nearly all the particles were localized within liposomes, confirming successful protection from the air-water interface (AWI). Importantly, particles were randomly distributed within the liposome rather than associating with the lipid bilayer, indicating minimal particle-lipid interactions. In contrast, control samples prepared using conventional graphene and holey carbon grids showed that most particle were adsorbed to the AWI or graphene surfaces. This interfacial adsorption is known to induce preferential orientation, complex disassembly, and denaturation ^4,5^.

**Figure 2.**
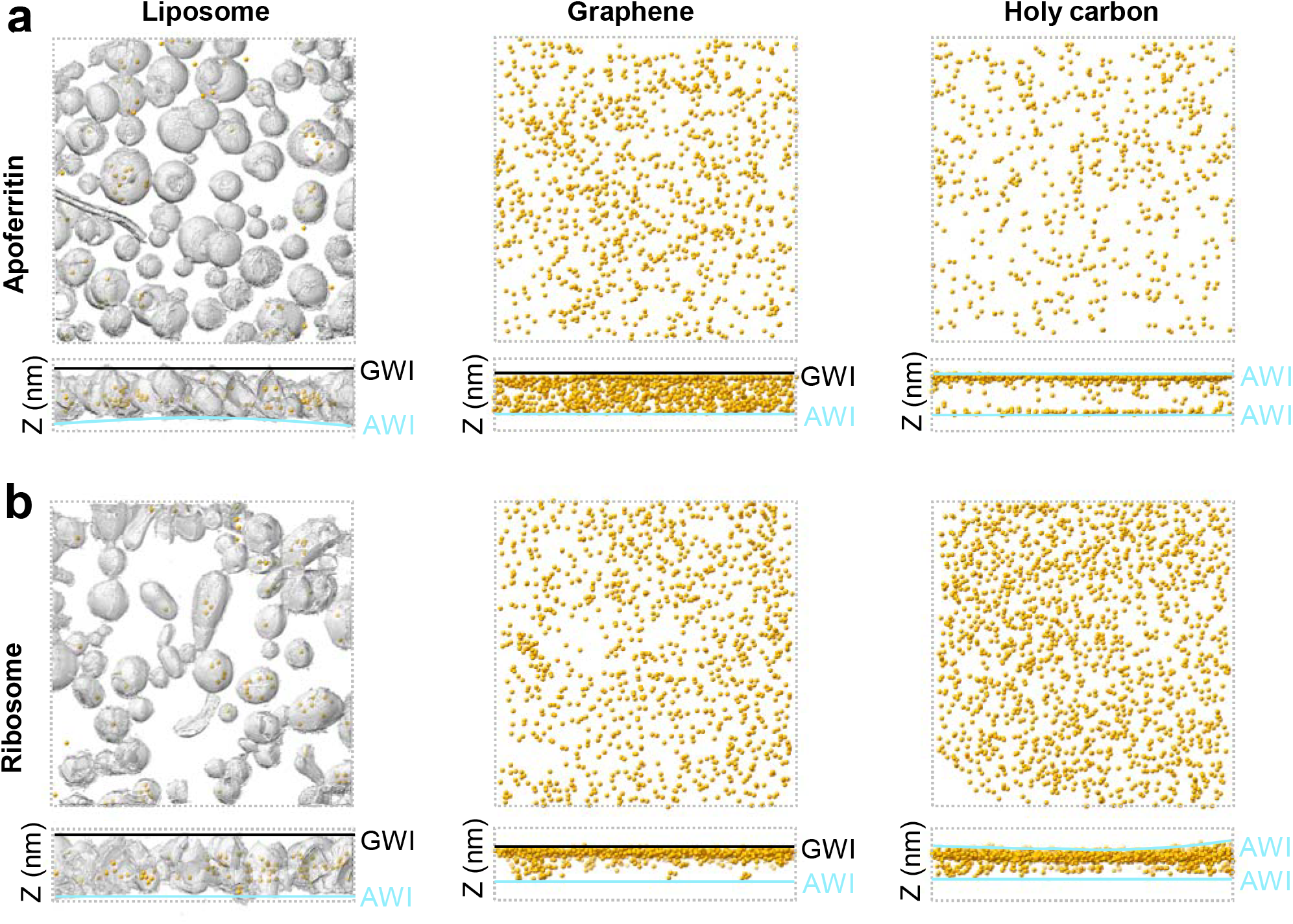
Distribution of the particles prepared with three methods. **a**, Representative tomograms of apoferritin sample prepared by liposome encapsulating (left panel), on graphene and holy carbon grid (middle and right panels). **b**, Representative tomograms of ribosome sample prepared by liposome encapsulating (left panel), on graphene and holy carbon grid (middle and right panels). The gray surface indicate liposome membrane, yellow dots indicate the position of macromolecules. Most of the particles were absorbed on the AWI or graphene-water interface.

### High-resolution structures could be resolved for the encapsulated samples

To further verify the liposome encapsulation method, we acquired cryo-EM datasets and performed single-particle analysis using CryoSPARC (v4.6.2) ^26^. Based on our encapsulation efficiency measurements (Figure S3), we prepared samples at high concentrations (50 μM for apoferritin, 14.2 μM for ribosome) to ensure sufficient particle density for high-resolution reconstruction. During data processing after data collection, 2D class averages revealed distinct populations: while some classes exhibited clear secondary structure features (Figure 3a, d, green boxes), others showed residual lipid membrane signals that compromised particle alignment (Figure 3a, d, red boxes). The latter likely represent either misaligned particles or false positives. To maximize particle retention, we included all 2D classes in subsequent masked 3D heterogeneous refinements. Following multiple rounds of refinement and CTF correction, we achieved final reconstructions at 3.32 Å resolution for apoferritin and 3.66 Å for ribosome (Figure 3b, e, and S5-S10), with well-resolved side-chain densities in both maps (Figure 3c, f).

**Figure 3.**
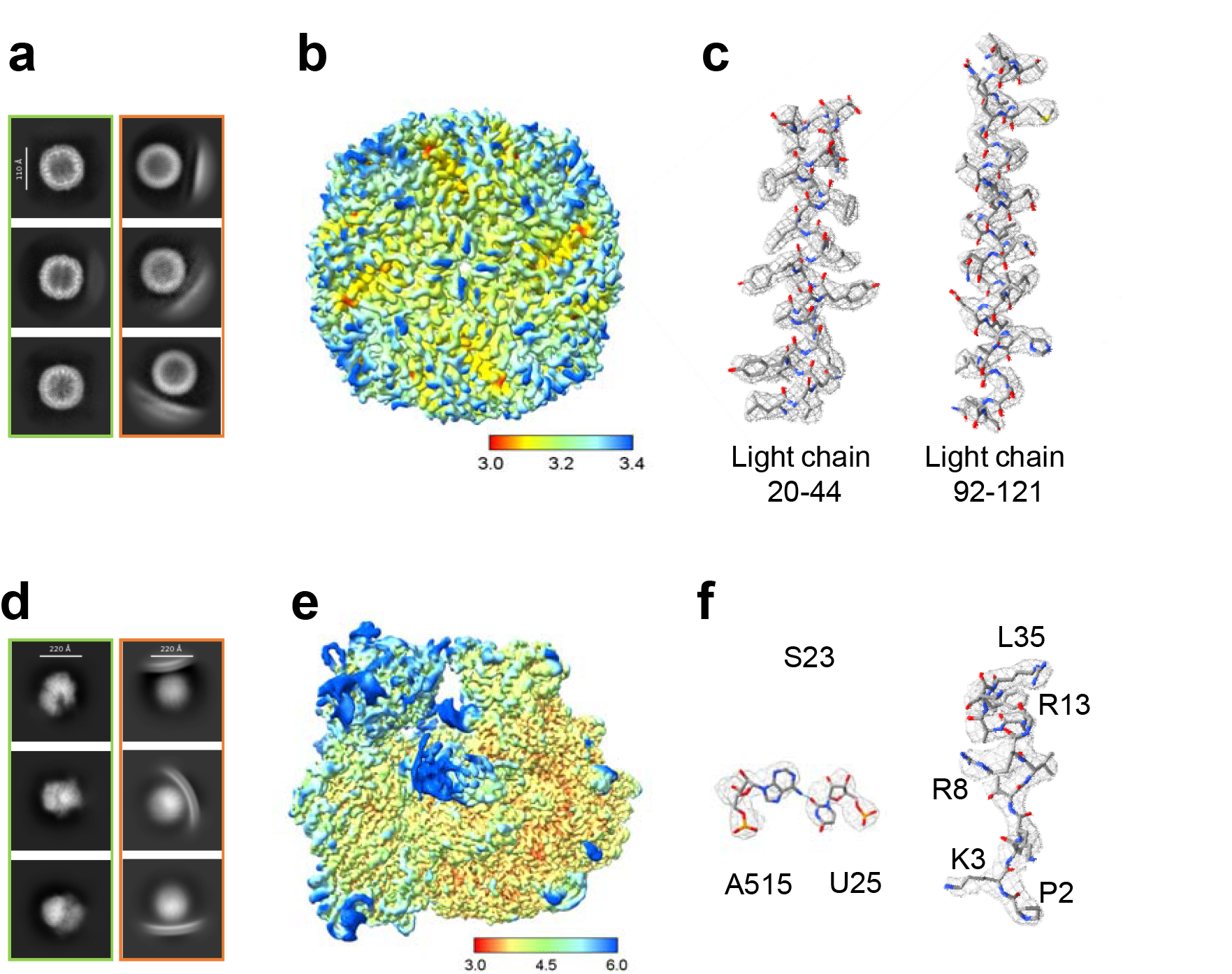
High resolution structures resolved for the liposome encapsulating sample. **a, d**, Representative 2D averages. 2D averages in the blue box are good aligned, while 2D averages in the red box are not well aligned because of signal from the lipid membrane. **b, e**, Local resolution map of both samples. **c, f**, Representative filtered density map from CryoSPARC overlapped with the refined atomic models (PDB ID: 2W0O, 5AFI).

To quantify and mitigate the impact of residual lipid signals on particle alignment, we systematically evaluated 2D masking strategies. Our analysis revealed that lipid interference became significant (>20% misalignment) when using masks exceeding 1.5× the particle diameter (∼12 nm) for apoferritin, compared to 1.4× (∼20 nm) for ribosome (Figure S11). These empirically determined thresholds provide practical guidelines for optimizing 2D classification in liposome-encapsulated samples.

### Assembling could be well protected in the liposome

The liposome encapsulation method provides a biomimetic environment that effectively shields macromolecules from the AWI, thereby solving common AWI-induced artifacts including complex dissociation, denaturation, and preferential particle orientation ^4,5^. To quantitatively assess this protective effect, we performed comparative single-particle analysis (using CryoSPARC v4.6.2 ^26^) between liposome-encapsulated samples and traditional preparations on graphene and holey carbon grids.

The *E. coli* 70S ribosome, comprising 50S and 30S subunits, maintains its structural integrity through divalent ions such as Mg^2+ 27^. This delicate equilibrium may be susceptible to AWI-induced destabilization. Indeed, our quantitative analysis revealed striking differences in ribosomal integrity: conventional preparations showed significant dissociation, with only 64% and 28% of particles kept assembled (consistent with previous report ^28^) on graphene-coated and holey carbon grids, respectively. In stark contrast, liposome-encapsulated samples showed no detectable sub-unit dissociation (Figure 4a), demonstrating the superior preservation capability of our method.

**Figure 4.**
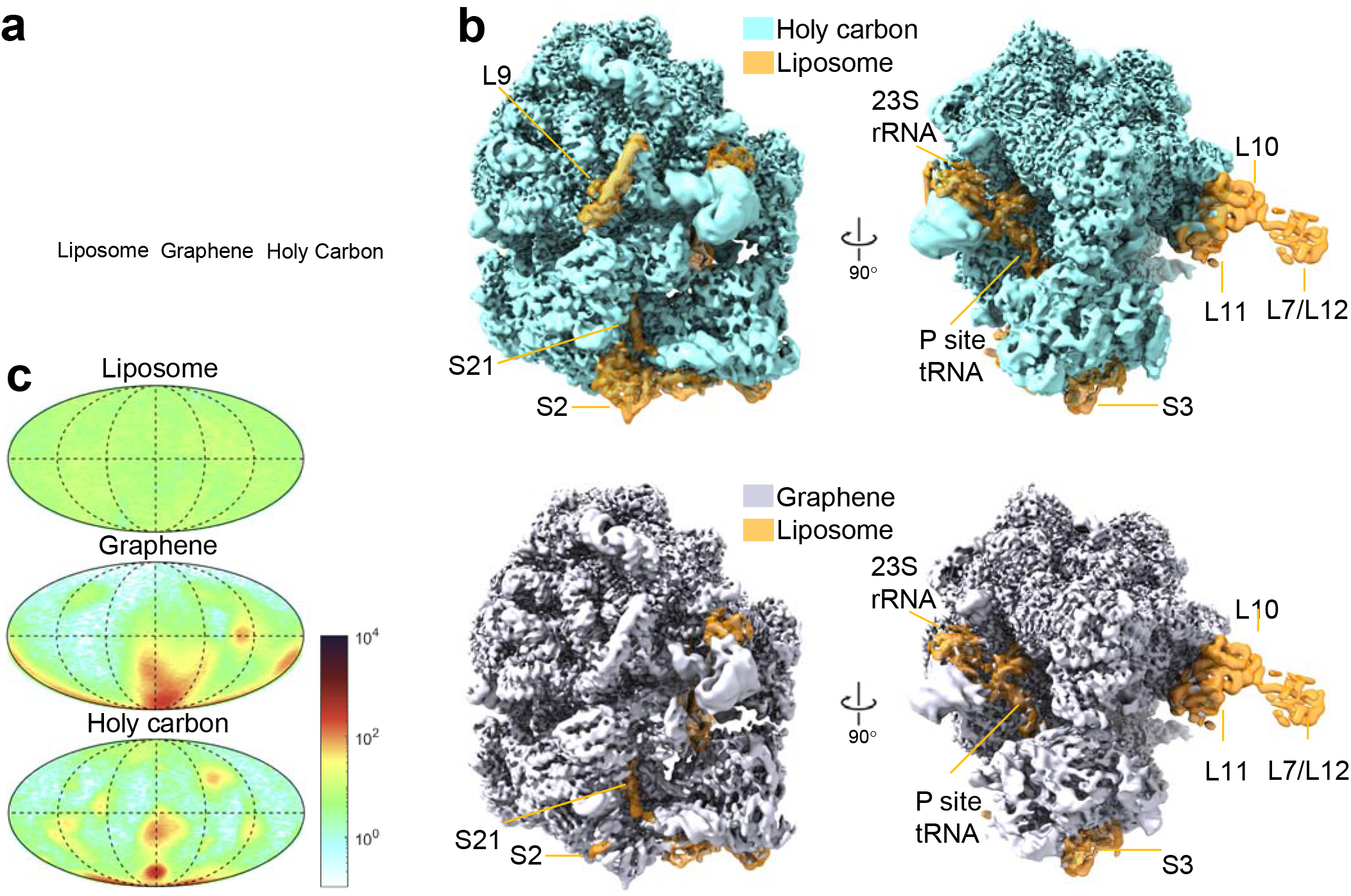
Structure analysis for the liposome encapsulating sample. **a**, Percentage of the assembled and disassembled 70s particles. The 50s particles was used to calculate the disassembled 70s particles. **b**, Comparsion of the 3D density maps of 70s particles. For the map reconstructed from the liposome encapsulated particles, only the density better than the control ones were highlighted. **c**, Mollweide projections of the 70s particles orientations.

### Surface structures could be well preserved in the liposome

Macromolecular surface is able to undergo local, transient unfolding ^7,9^. The AWI may perturb the folding-unfolding equilibrium ^3^, which is potentially inducing structural artifacts. However, the liposome encapsulation method may protect against such interfacial disturbances by providing a native-like microenvironment. Comparative structural analysis revealed distinct preservation advantages in liposome-encapsulated ribosomes versus traditional preparations (on graphene and holey carbon grid). Besides the densities that have reserved in the control maps, the liposome-derived reconstruction showed significantly improved density at the surface-exposed components including: Protein sub-units: L7/L12, L9, L10, L11, S2, S3, S21; RNA elements: 23S rRNA; and functional sites: P-site tRNA (Figure 4b). Notably, the most dramatic preservation improvement occurred in the L7/L12 stalk complex (including adjacent L10 and L11 proteins), which is known a critical functional domain. The observed pattern of structural preservation — where only specific sub-regions showed reduced density rather than complete subunit loss — suggests our method effectively minimizes AWI-induced denaturation while maintaining overall particle integrity.

### Particle orientation could be well distributed in the liposome

Previous studies have demonstrated that 70S ribosome particles exhibit strong preferential orientation on conventional cryo-EM grids, with alleviation observed when using an ultra-thin carbon support ^29^. Our control experiments reproduced this finding, confirming persistent orientation bias even with graphene-modified grids (Figure 4c). In striking contrast, liposome-encapsulated ribosomes showed near-ideal isotropic angular distribution (Figure 4c), effectively resolving the long-standing orientation preference problem in cryo-EM. The orientation behavior of apoferritin is different due to its high symmetry (octahedral point group). As expected for such symmetric particles, all preparation methods — including conventional grids and our liposome approach — yielded essentially uniform angular distributions (Figures S5-S7). Altogether, the orientation benefits of liposome encapsulation are particularly valuable for asymmetric complexes like ribosomes.

## Discussion

A key advantage of cryo-EM over X-ray crystallography is the potential to study macromolecules in a more near-native aqueous environments. However, conventional sample preparation on holey carbon grids results in >90% of particles adsorbing to the AWI, where the altered folding/unfolding energy landscape ^3^ may yield non-native conformations or artificial structural features. More critically, AWI exposure frequently induces preferential orientation, complex disassembly, or complete denaturation, making high-resolution structure determination inefficient or impossible. To address these limitations, we developed a liposome encapsulation approach combined with graphene support grids. This method physically confines macromolecules within lipid vesicles, maintaining a solution-like environment throughout the whole process of specimen preparation. Cryo-electron tomography confirmed successful encapsulation of both apoferritin and 70S ribosomes, with particles randomly distributed within liposomes and completely isolated from the AWI (Figure 2a). The absence of particle adsorption to liposome membranes — consistent with previous reports of non-fouling surfaces ^11,24,25^ — explains the dramatic improvements in angular distribution, surface feature preservation, and complex integrity observed in our reconstructions (Figure 3, 4).

We validated this method using two model systems spanning a wide mass range (0.443-2.3 MDa). Here we chose the liposome size being 100 nm, which is larger than most samples used in cryo-EM. While 100 nm liposomes accommodated these large complexes effectively, the current implementation may face challenges with smaller (<200 kDa) macromolecules due to signal dilution in the relatively thick ice layers required. To study smaller sized macromolecules, smaller sized liposomes such as ∼50 nm in diameter could be used. Unlike conventional blotting methods that require extensive optimization of sample-dependent parameters (concentration, blotting conditions, etc.), our approach uses a standardized preparation protocol for all water-soluble macromolecules. This intrinsic sample-independence significantly improves reproducibility and accessibility. We anticipate this method will enable high-resolution structural studies of numerous challenging complexes that are currently intractable due to AWI-induced problems.

### Methods Protein sources

The apoferritin (from equine spleen) was bought from Sigma-Aldrich (CAS NO. A3660). The *E*.*coli* ribosome was bought from NEB (CAS NO. P0763S).

### Liposome encapsulated sample preparation

The phospholipids used to prepare liposomes were 1,2-dioleoyl-sn-glycero-3 phosphocholine (DOPC, Avanti Polar Lipids) and 1,2-dioleoyl-sn-glycero-3-phospho-L-serine (DOPS, Avanti Polar Lipids). In addition, a small fraction of 1,2-dipalmitoyl-sn-glycero-3-phosphoethanolamine-N-(cap Cy5) (Cy5-cap PE, Avanti Polar Lipids) could be added to track position and concentration of the lipid. A series of lipid ratio were screened for best result. Finally, 95% DOPC, 5% DOPS were used to make liposomes. The lipid mixture was first dissolved in chloroform in a glass vial (1-2 mL). Chloroform was then removed by continuous mild nitrogen gas flow. To completely remove the chloroform and have dry lipid cake, the vial was vacuumed for 6h with a water pump. Later, the sample was added to the glass vial and shacked gently to resolve the lipid cake to form liposomes. Then, the sample was forced to go through a polycarbonate filter with 100 nm pore size (GE Healthcare) using a mini-extruder (Avanti Polar Lipids) for 201 times in order to get the final liposomes with proteins packed in. To minimize to sample volume, we modified the Teflon block in the extruder and made the dead volume 10 times smaller (10 μL). By this way, much smaller volume (30-50 μL) of sample could be prepared. This volume is similar to the common smallest dead volume (30-40 uL) of the ultrafiltration tube.

### Unencapsulated sample digestion

The macromolecules that outside the liposomes are not wanted and should be removed. The unpacked apoferritin liposomes were digested with proteinase K (Sangon Biotech, CAS No. B600169, same mass fraction as apoferritin) at 37 °C and then ultrafiltrated. Apoferritin is very stable, so we repeated digestion for four times to completely remove the unpacked apoferritin particles: with 3h for the first time, 6h for the second time, 12h for the third time, and 24h for the last time. Between two digestions, the sample was added to the ultrafiltration tube (100,000 MWCO, Vivaspin Turbo 4, Sartorius) and centrifuged at 1000-2000 × g to remove the digested fragments. For the ribosome, the unpacked ribosomes were digested with proteinase K (Sangon Biotech, CAS No. B600169) and OmniNuclease (Sangon Biotech, CAS No. B610168) for 5h at 37 °C. Finally, the samples were concentrated to about 20 mg/ml (lipid concentration) for cryo-EM sample preparation.

### Cryo-EM sample preparation

The samples (3.5 μL) were applied to pre-glow-discharged grids. Several types of grids were tested, including homemade graphene grid ^8,15^ based on a holy carbon grid (300 mesh, QUANTIFOIL® R 1.2/1.3); homemade graphene oxide (GO) grid ^30^ based on a holy carbon grid (300 mesh, QUANTIFOIL® R 1.2/1.3); Lacey carbon film with a continuous layer of ultrathin carbon film (300 mesh, QUANTIFOIL® R 1.2/1.3); Holy carbon grid (300 mesh, QUANTIFOIL® R 1.2/1.3). The grids were blotted for around 4 seconds at 100% relative humidity, wait for 30s and then plunge-frozen in liquid ethane using a FEI Mark IV Vitrobot.

### Cryo-EM imaging and data collection

For the initial screening, we loaded cryo-grids into a Thermo Scientific™ Glacios™ transmission electron microscope, which was operated at 200 kV. Then, the grids with good ice thickness were used for data collection. Single particle data were collected using EPU on a Titan Krios transmission electron microscope operated at 300 kV, equipped with K3 detectors and energy filter. The respective imaging pixel sizes used in the data collection were 1.04 Å. The defocus ranged from -0.5 to -2.5 and the total dose was 50 electrons per Å^2^ which is divided into 40 frames (Supplementary Table 1).

### Cryo-EM data processing and 3D reconstruction

Data processing was performed using CryoSPARC (v4.6.2) ^26^. Following motion correction and patch CTF extraction of the movie frames, particles were selected by blob picking. Particles were then extracted for multiple rounds of 2D classification. The particles from good 2D classes that showed particle features were retained for further homogeneous and heterogeneous refinement. Good maps with better overall structure features were selected for further homo-refinement, global and local CTF refinement ^31^ During ribosome reconstruction, to calculate the number of disassembled 50s and 30s, correct 3D volumes of 70s, 50s and 30s were used as references in the heterogeneous refinement jobs. Reconstruction information is presented in Supplementary Table 1, and image processing and 3D reconstruction steps are depicted in Figure S5-S10.

### Cryo-ET data collection

Cryo-ET tilt series were acquired using a Thermo Fisher Titan Krios operated at 300□keV equipped with Falcon 4 direct electron detector in Shuimu BioSciences Ltd. Tilt series were collected with SerialEM ^32^ with a physical pixel size of 2.2□Å per pixel. They were acquired using dose-symmetric tilt-scheme starting from 0° with a 3° tilt increment by a group of two and an angular range of ±51°. The accumulated dose of each tilt series was around 122.5 e^-^/Å^2^ with a defocus range between −3 and −5 μm. Eight frames were saved in each raw tilt image. Details of data collection parameters are listed in Supplementary Table 2.

### Cryo-ET data processing and 3D reconstruction

Video frames of each tilt were motion-corrected, summed and binned with a factor of two by using MotionCor2 ^33^. Tilt series were merged into one stack and aligned by AreTomo ^34^. Tomograms were three-dimensionally reconstructed by the SIRT (Simultaneous Iterative Reconstruction Technique) method. 3D visualization was made by IMOD v.4.10.38 ^35^, Chimera v.1.14 ^36^ and ChimeraX v.1.4 ^37^. Particle picking were done by emClarity ^38^. Liposome segmentation was by MemBrain-Seg ^39^.

## Data availability

All data needed to evaluate the conclusions in the paper are present in the paper and/or the Supplementary information and source data are provided with this paper. The cryo-EM density maps and corresponding atomic models have been deposited in the EMDB and PDB, respectively. The accession codes are: XXXXXX.

## Acknowledgement

This work was supported by National Natural Science Foundation of China [No. T2221001 to M.L., and No. 32371286 to J.M.]; the Strategic Priority Research Program of the Chinese Academy of Sciences (XDB0480000); the Chinese Academy of Sciences Project for Young Scientists in Basic Research (YSBR-104); National Key Research and Development Program of China [No. 2019YFA0709304 to M.L.]; and the start-up grants of Wenzhou Institute, University of Chinese Academy of Sciences (WIUCASQD2025069). We thank Beijing National Laboratory for Condensed Matter Physics Institute of Physics Chinese Academy of Science and Beijing Branch of Songshan Lake Laboratory for Materials Science for support of cryo-EM data acquisition. We also thank Zhenqian Guo for his assistance in cryo-ET data acquisition in the Shuimu BioSciences Ltd.

## Author contributions

XXXXXX.

