## Supplemental Information for "Liposome Encapsulation Enables Near-Native Cryo-EM Structural Determination by Shielding Macromolecules from Non-Physiological Interfaces"

### Slide 1
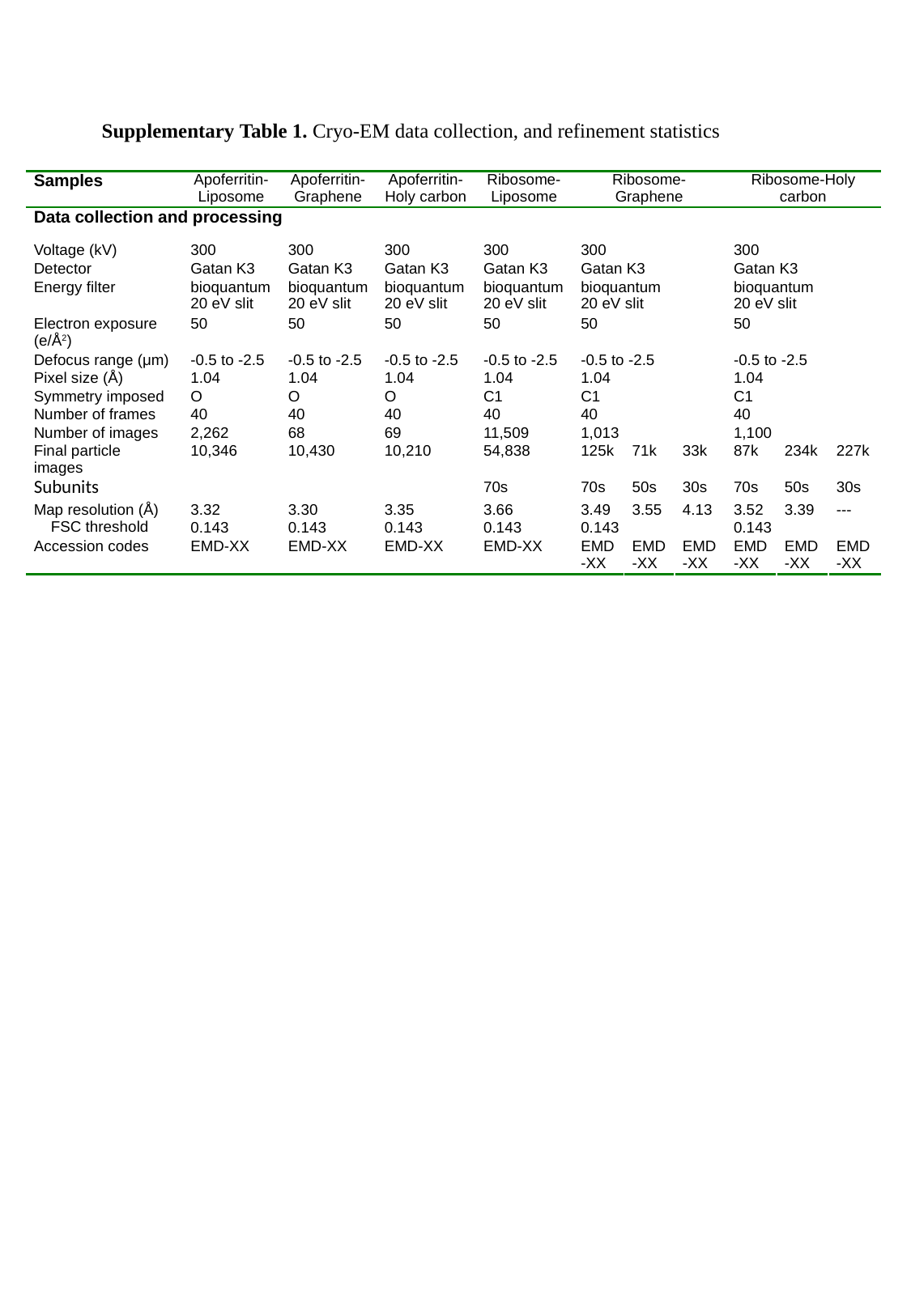

Supplementary Table 1. Cryo-EM data collection, and refinement statistics
| Samples | Apoferritin-Liposome | Apoferritin-Graphene | Apoferritin-Holy carbon | Ribosome-Liposome | Ribosome-Graphene | | | Ribosome-Holy carbon | | |
| --- | --- | --- | --- | --- | --- | --- | --- | --- | --- | --- |
| Data collection and processing | | | | | | | | | | |
| Voltage (kV) | 300 | 300 | 300 | 300 | 300 | | | 300 | | |
| Detector | Gatan K3 | Gatan K3 | Gatan K3 | Gatan K3 | Gatan K3 | | | Gatan K3 | | |
| Energy filter | bioquantum 20 eV slit | bioquantum 20 eV slit | bioquantum 20 eV slit | bioquantum 20 eV slit | bioquantum 20 eV slit | | | bioquantum 20 eV slit | | |
| Electron exposure (e/Å^2^) | 50 | 50 | 50 | 50 | 50 | | | 50 | | |
| Defocus range (μm) | -0.5 to -2.5 | -0.5 to -2.5 | -0.5 to -2.5 | -0.5 to -2.5 | -0.5 to -2.5 | | | -0.5 to -2.5 | | |
| Pixel size (Å) | 1.04 | 1.04 | 1.04 | 1.04 | 1.04 | | | 1.04 | | |
| Symmetry imposed | O | O | O | C1 | C1 | | | C1 | | |
| Number of frames | 40 | 40 | 40 | 40 | 40 | | | 40 | | |
| Number of images | 2,262 | 68 | 69 | 11,509 | 1,013 | | | 1,100 | | |
| Final particle images | 10,346 | 10,430 | 10,210 | 54,838 | 125k | 71k | 33k | 87k | 234k | 227k |
| Subunits | | | | 70s | 70s | 50s | 30s | 70s | 50s | 30s |
| Map resolution (Å) FSC threshold | 3.32 | 3.30 | 3.35 | 3.66 | 3.49 | 3.55 | 4.13 | 3.52 | 3.39 | --- |
| | 0.143 | 0.143 | 0.143 | 0.143 | 0.143 | | | 0.143 | | |
| Accession codes | EMD-XX | EMD-XX | EMD-XX | EMD-XX | EMD-XX | EMD-XX | EMD-XX | EMD-XX | EMD-XX | EMD-XX |

### Slide 2
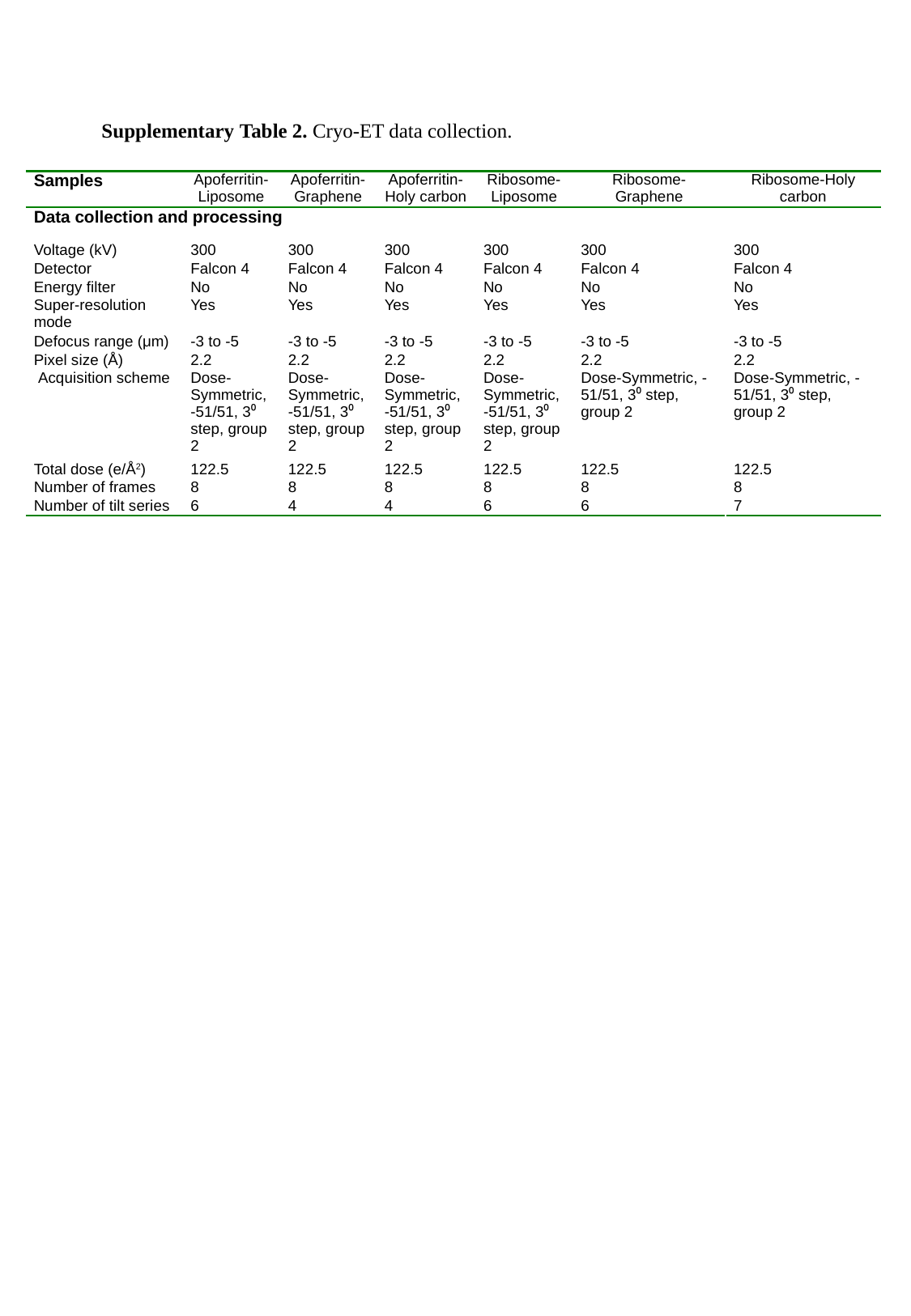

Supplementary Table 2. Cryo-ET data collection.
| Samples | Apoferritin-Liposome | Apoferritin-Graphene | Apoferritin-Holy carbon | Ribosome-Liposome | Ribosome-Graphene | Ribosome-Holy carbon |
| --- | --- | --- | --- | --- | --- | --- |
| Data collection and processing | | | | | | |
| Voltage (kV) | 300 | 300 | 300 | 300 | 300 | 300 |
| Detector | Falcon 4 | Falcon 4 | Falcon 4 | Falcon 4 | Falcon 4 | Falcon 4 |
| Energy filter | No | No | No | No | No | No |
| Super-resolution mode | Yes | Yes | Yes | Yes | Yes | Yes |
| Defocus range (μm) | -3 to -5 | -3 to -5 | -3 to -5 | -3 to -5 | -3 to -5 | -3 to -5 |
| Pixel size (Å) | 2.2 | 2.2 | 2.2 | 2.2 | 2.2 | 2.2 |
| Acquisition scheme | Dose-Symmetric, -51/51, 3⁰ step, group 2 | Dose-Symmetric, -51/51, 3⁰ step, group 2 | Dose-Symmetric, -51/51, 3⁰ step, group 2 | Dose-Symmetric, -51/51, 3⁰ step, group 2 | Dose-Symmetric, -51/51, 3⁰ step, group 2 | Dose-Symmetric, -51/51, 3⁰ step, group 2 |
| Total dose (e/Å^2^) | 122.5 | 122.5 | 122.5 | 122.5 | 122.5 | 122.5 |
| Number of frames | 8 | 8 | 8 | 8 | 8 | 8 |
| Number of tilt series | 6 | 4 | 4 | 6 | 6 | 7 |

### Slide 3
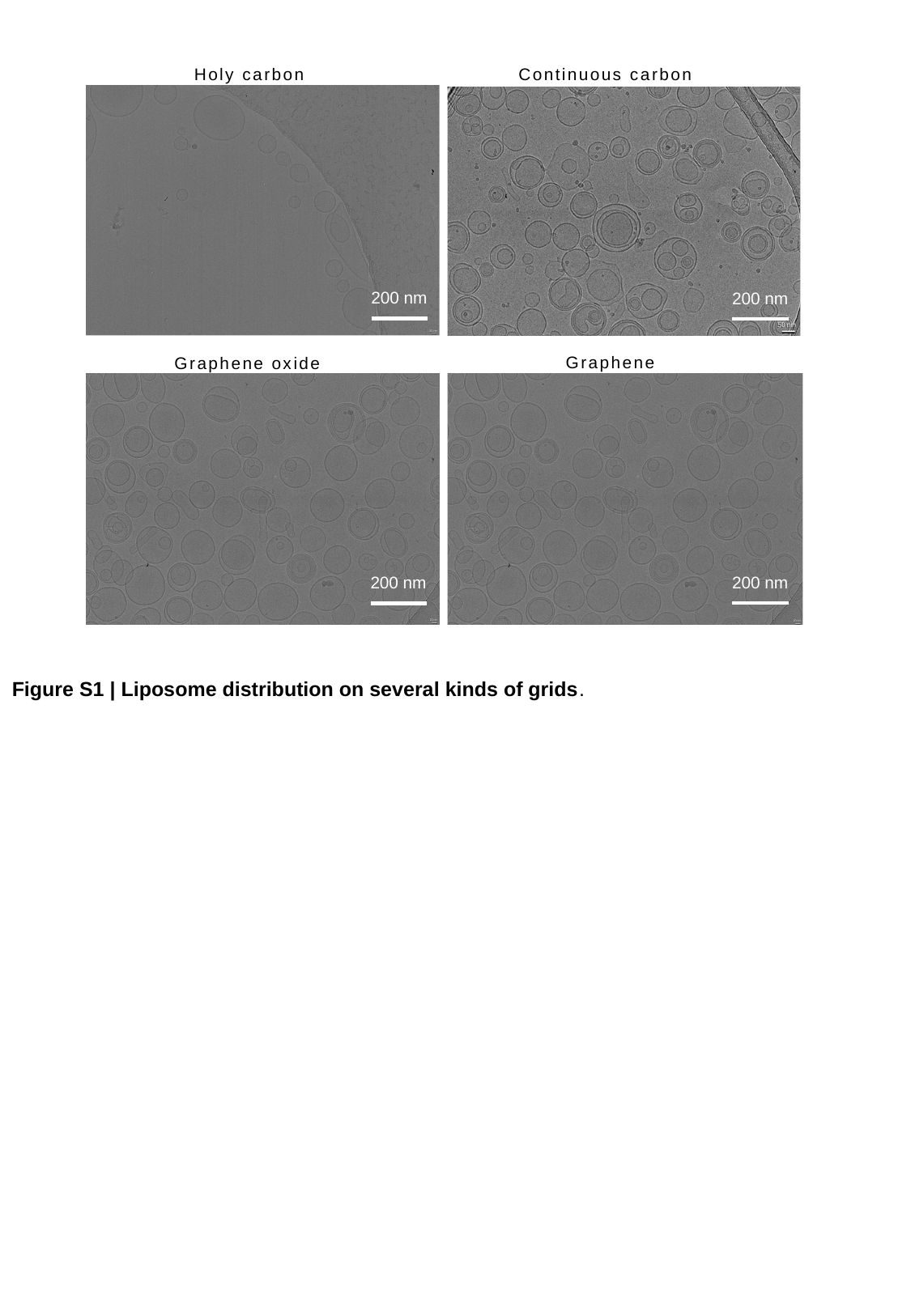

Holy carbon
Continuous carbon
200 nm
200 nm
Graphene
Graphene oxide
100 nm
200 nm
200 nm
Figure S1 | Liposome distribution on several kinds of grids.

### Slide 4
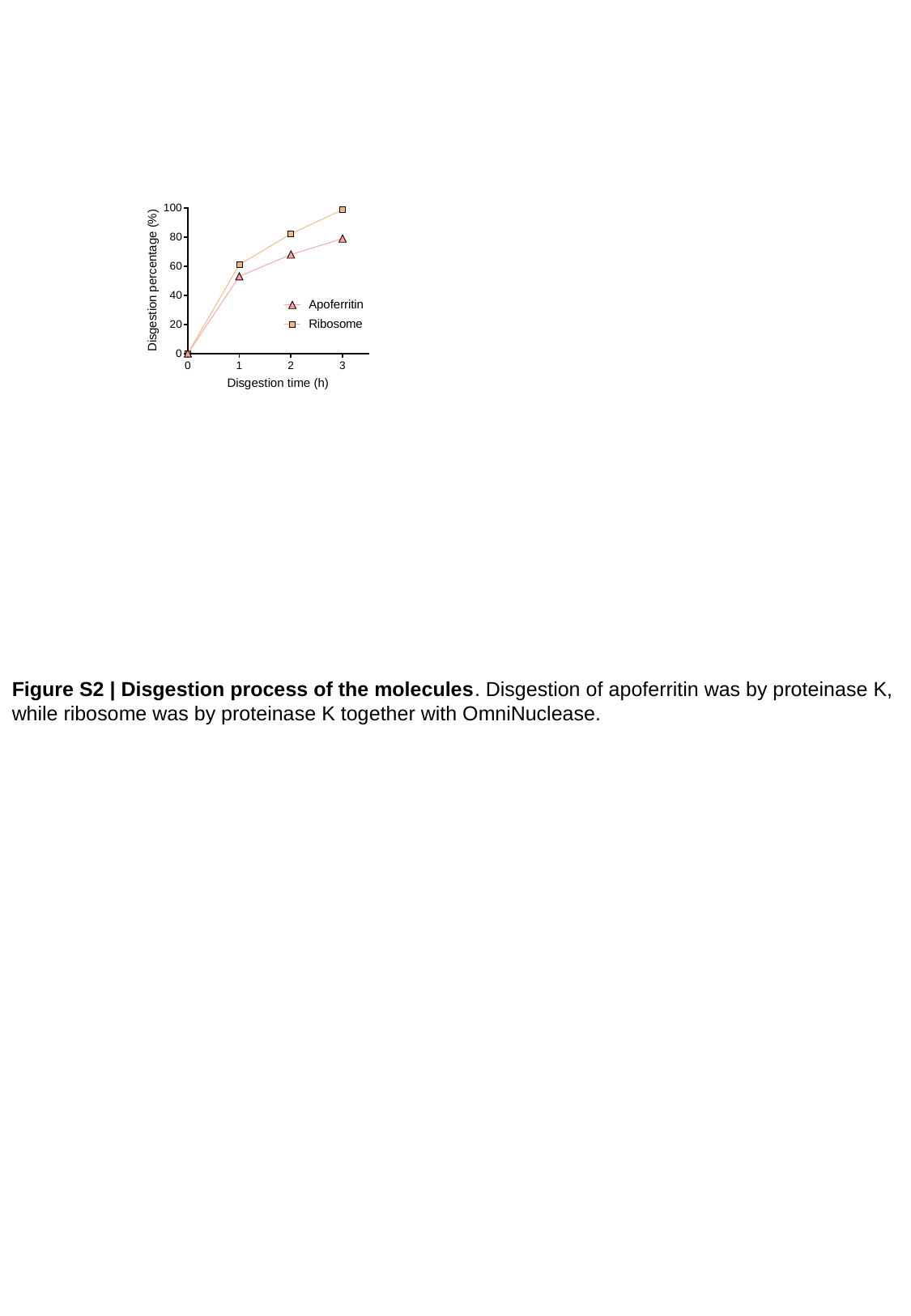

Figure S2 | Disgestion process of the molecules. Disgestion of apoferritin was by proteinase K, while ribosome was by proteinase K together with OmniNuclease.

### Slide 5
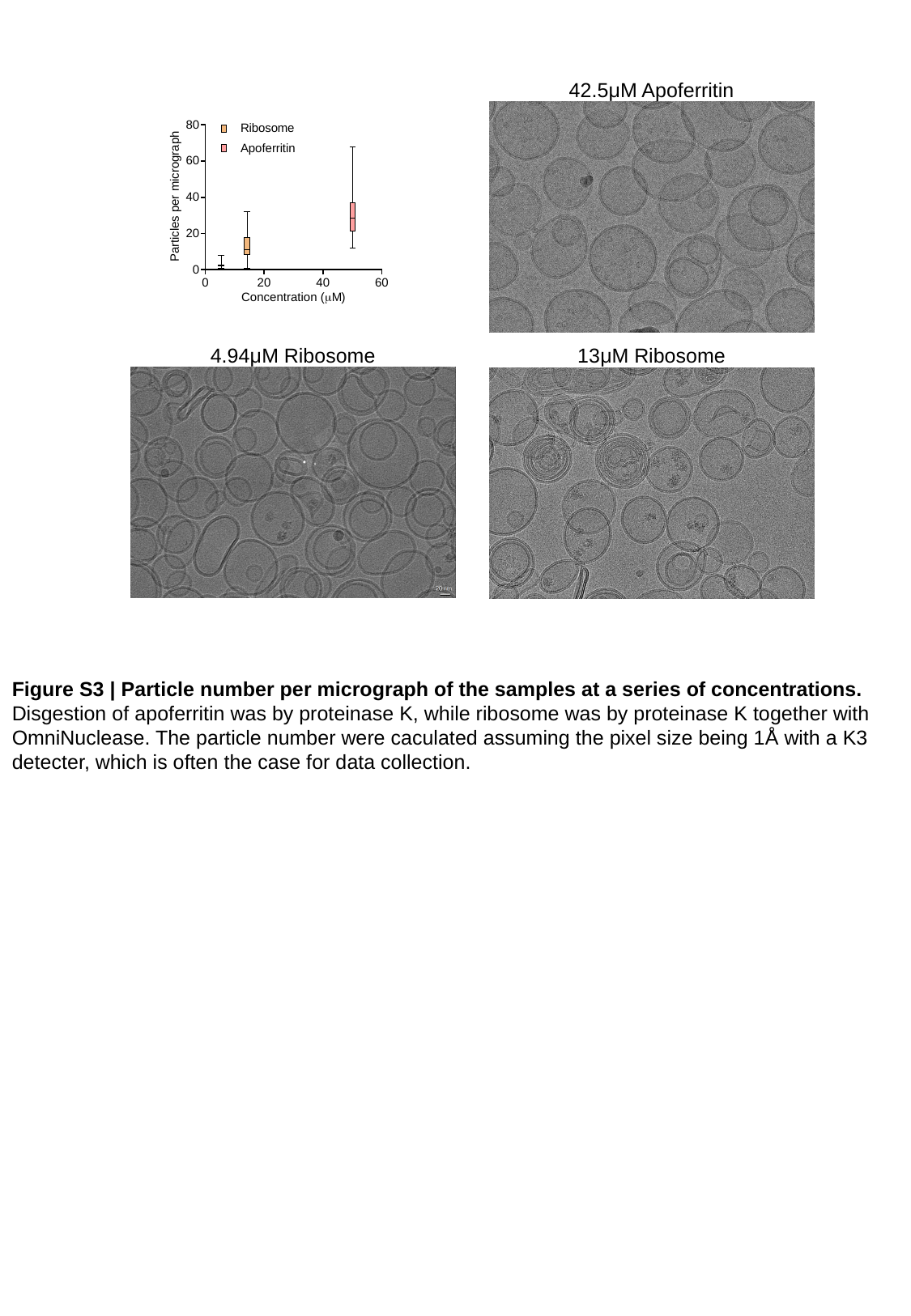

42.5μM Apoferritin
4.94μM Ribosome
13μM Ribosome
Figure S3 | Particle number per micrograph of the samples at a series of concentrations. Disgestion of apoferritin was by proteinase K, while ribosome was by proteinase K together with OmniNuclease. The particle number were caculated assuming the pixel size being 1Å with a K3 detecter, which is often the case for data collection.

### Slide 6
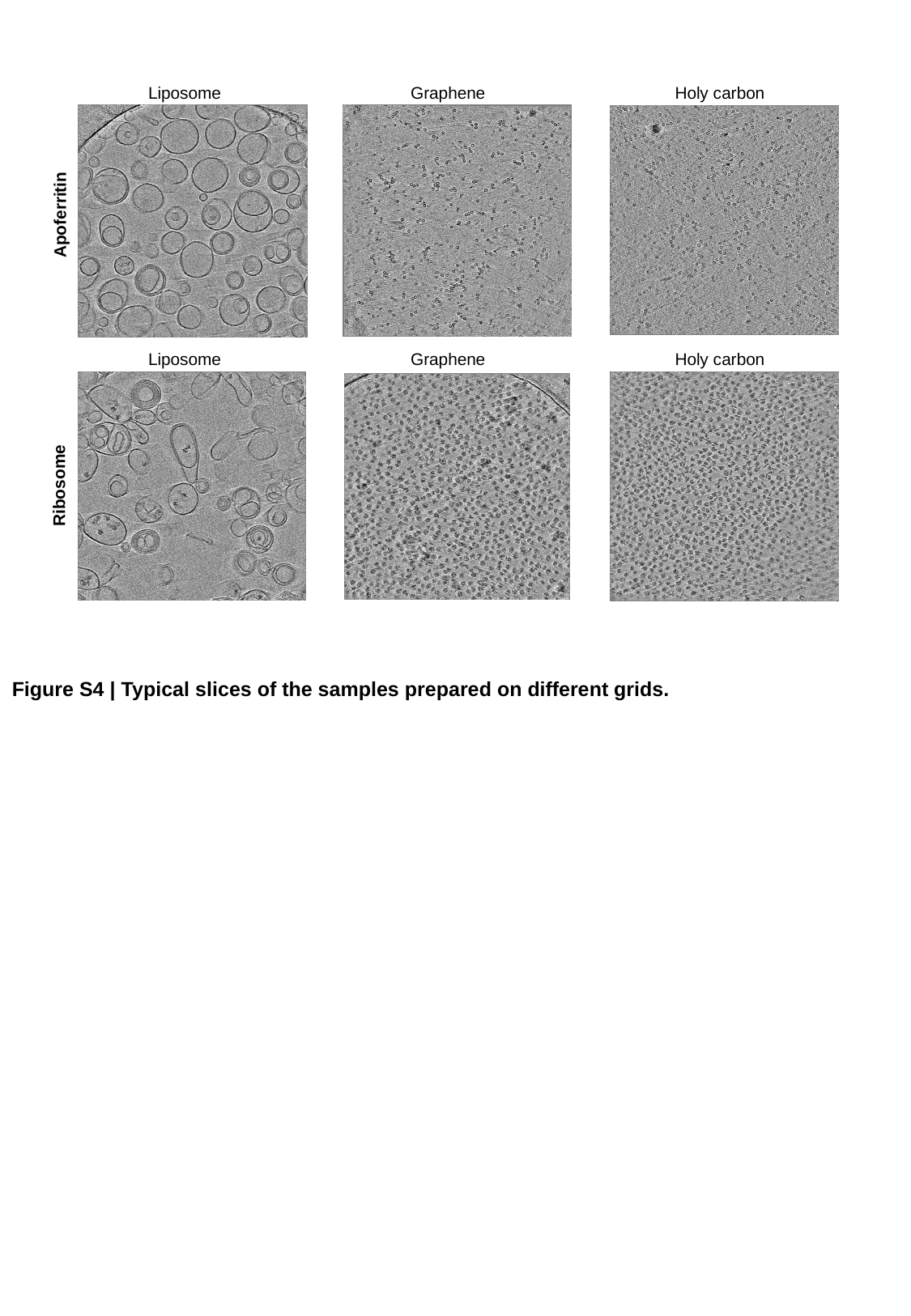

Liposome
Graphene
Holy carbon
Apoferritin
Liposome
Graphene
Holy carbon
Ribosome
Figure S4 | Typical slices of the samples prepared on different grids.

### Slide 7
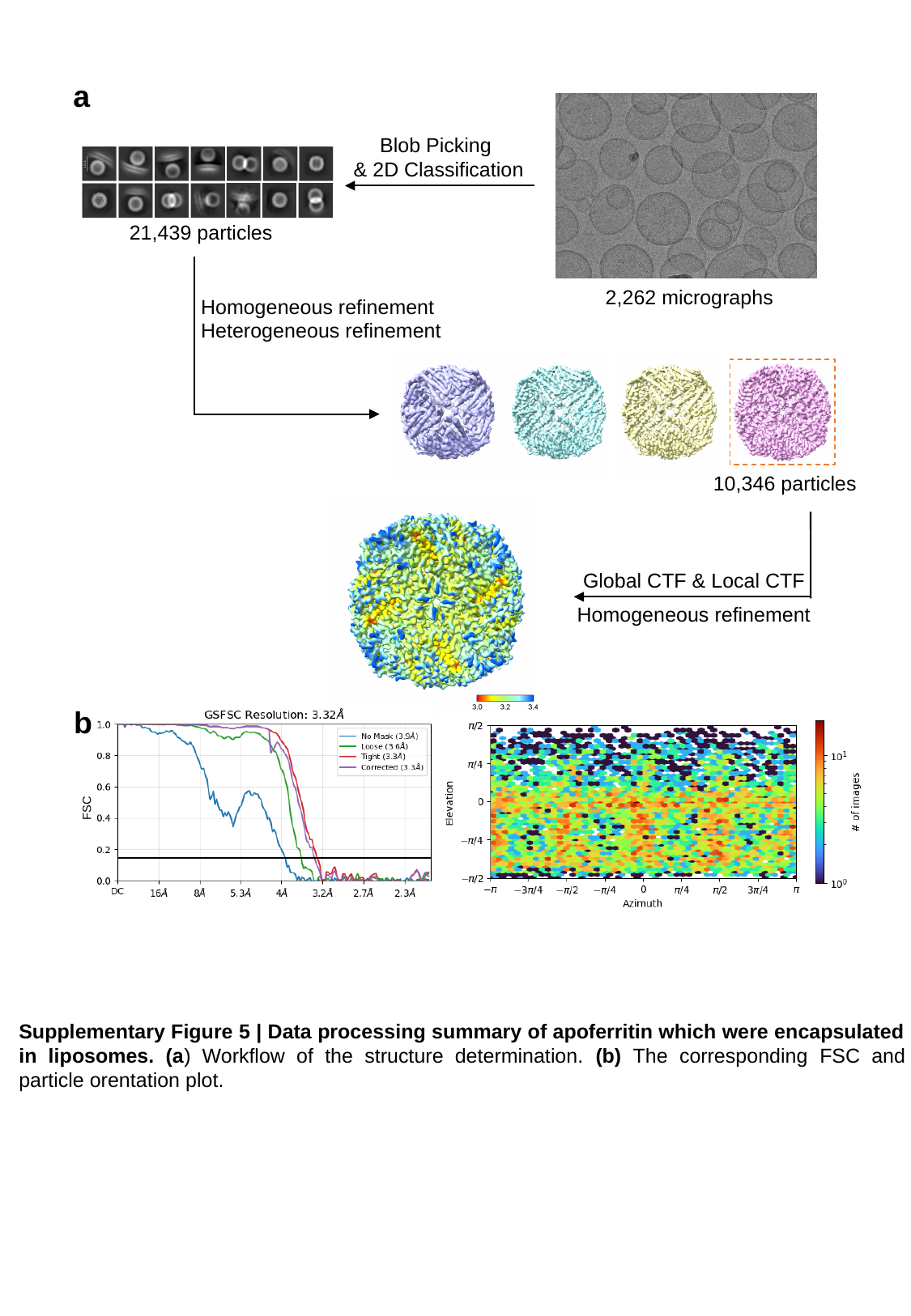

a
Blob Picking
 & 2D Classification
21,439 particles
2,262 micrographs
Homogeneous refinement
Heterogeneous refinement
10,346 particles
Global CTF & Local CTF
Homogeneous refinement
b
FSC
Supplementary Figure 5 | Data processing summary of apoferritin which were encapsulated in liposomes. (a) Workflow of the structure determination. (b) The corresponding FSC and particle orentation plot.

### Slide 8
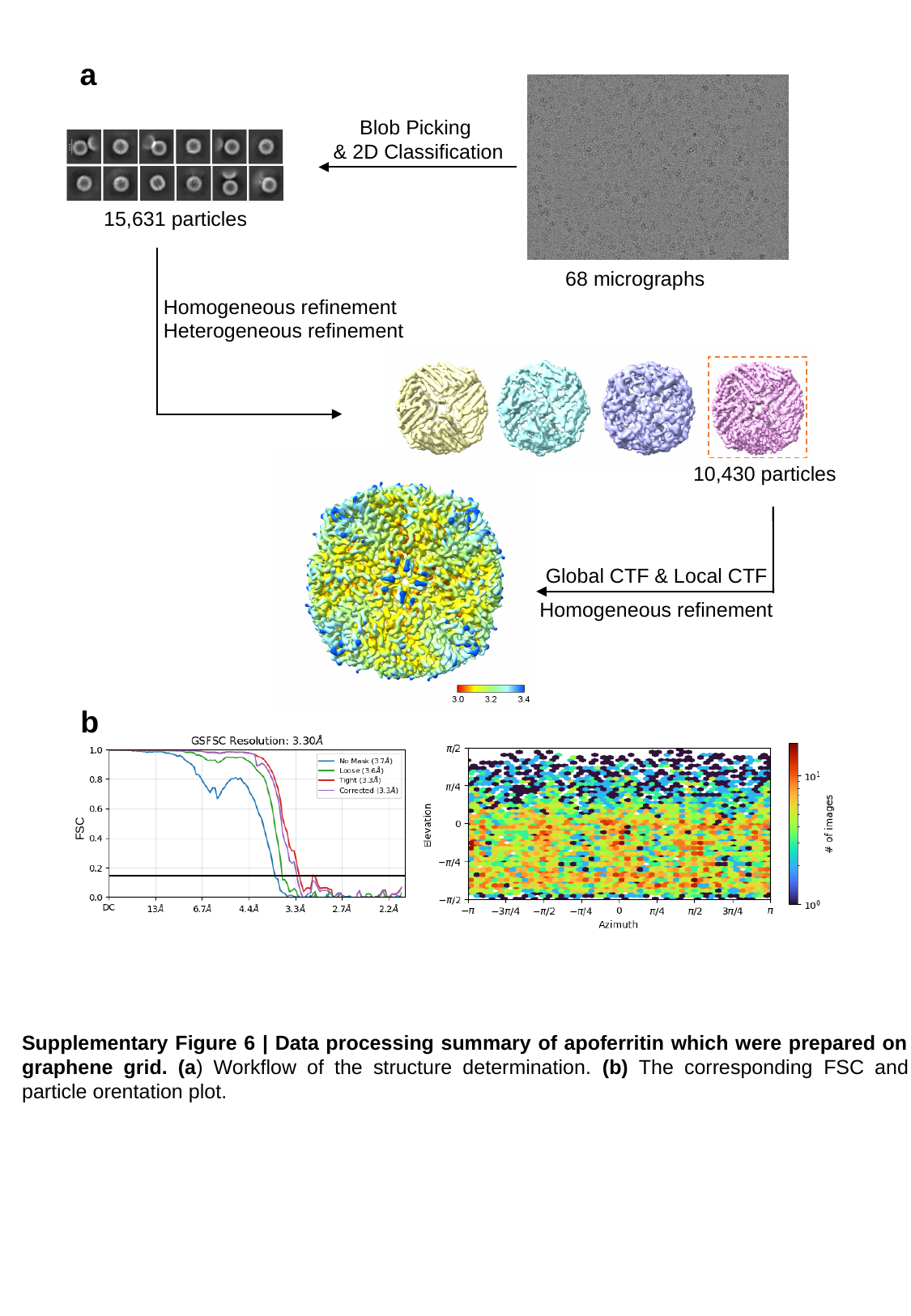

a
Blob Picking
 & 2D Classification
15,631 particles
68 micrographs
Homogeneous refinement
Heterogeneous refinement
10,430 particles
Global CTF & Local CTF
Homogeneous refinement
b
FSC
Supplementary Figure 6 | Data processing summary of apoferritin which were prepared on graphene grid. (a) Workflow of the structure determination. (b) The corresponding FSC and particle orentation plot.

### Slide 9
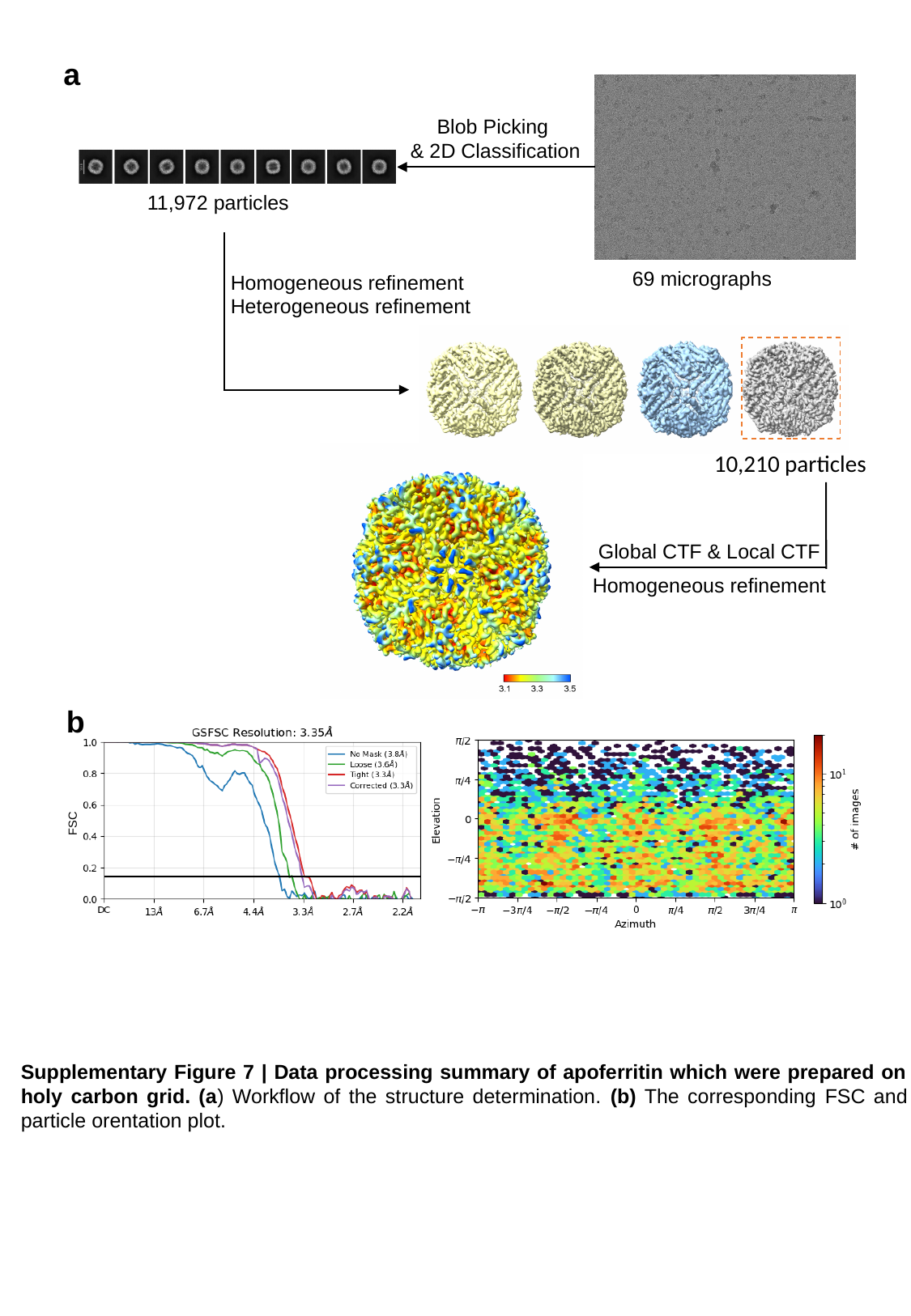

a
Blob Picking
 & 2D Classification
11,972 particles
69 micrographs
Homogeneous refinement
Heterogeneous refinement
10,210 particles
Global CTF & Local CTF
Homogeneous refinement
b
FSC
Supplementary Figure 7 | Data processing summary of apoferritin which were prepared on holy carbon grid. (a) Workflow of the structure determination. (b) The corresponding FSC and particle orentation plot.

### Slide 10
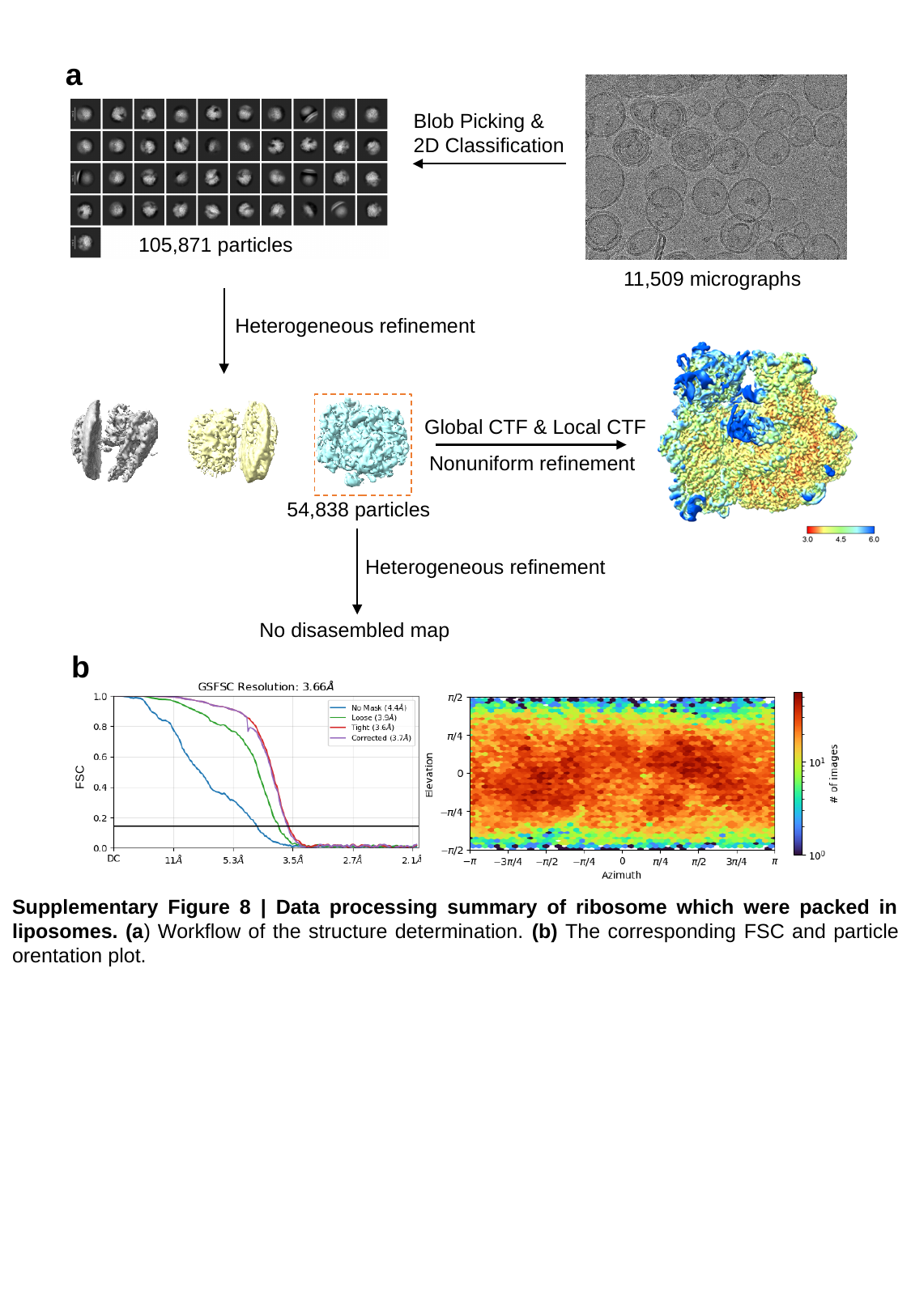

a
Blob Picking &
2D Classification
105,871 particles
11,509 micrographs
Heterogeneous refinement
Global CTF & Local CTF
Nonuniform refinement
54,838 particles
Heterogeneous refinement
No disasembled map
b
FSC
Supplementary Figure 8 | Data processing summary of ribosome which were packed in liposomes. (a) Workflow of the structure determination. (b) The corresponding FSC and particle orentation plot.

### Slide 11
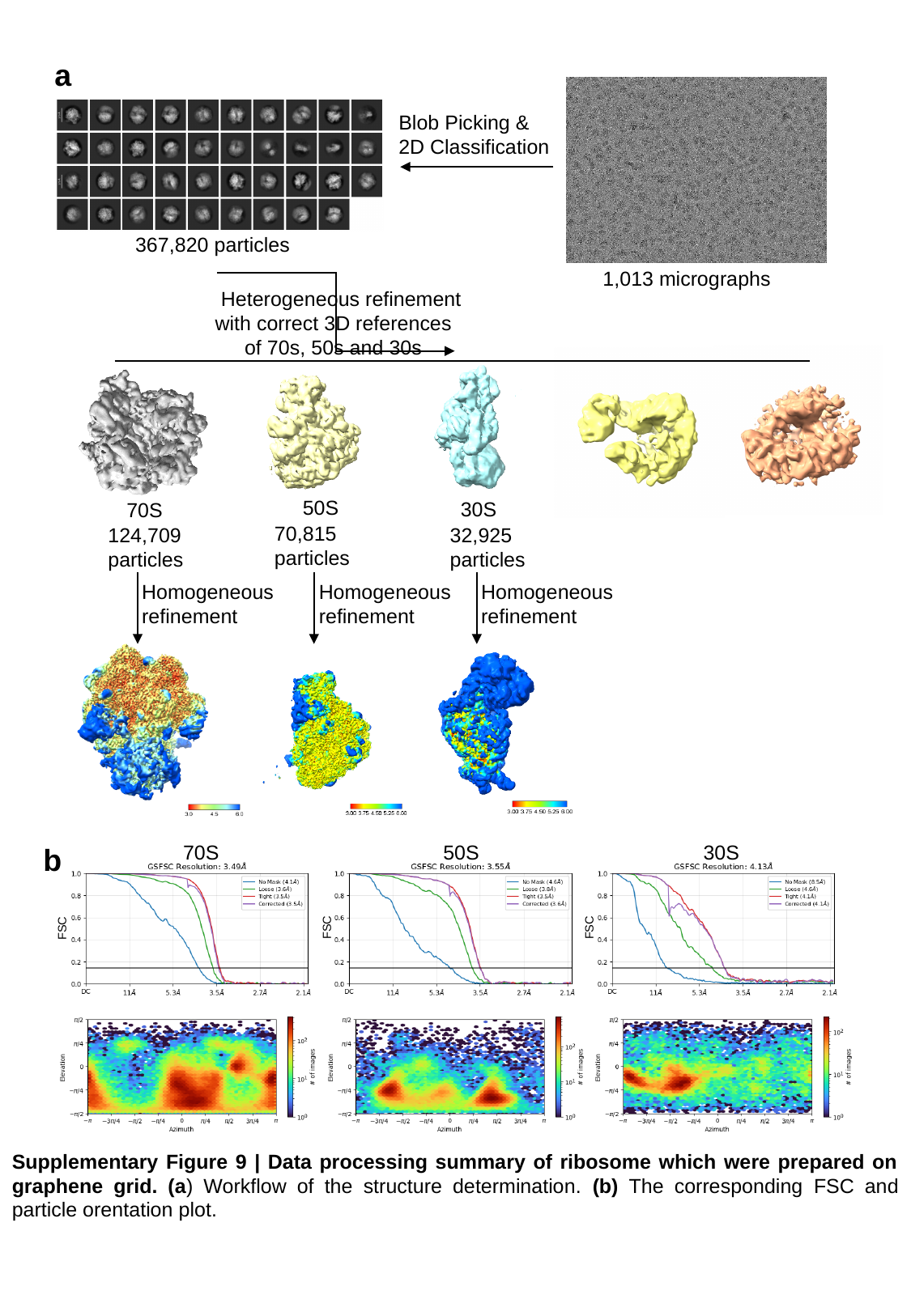

a
Blob Picking &
2D Classification
367,820 particles
1,013 micrographs
Heterogeneous refinement
with correct 3D references of 70s, 50s and 30s
50S
30S
70S
70,815
particles
124,709 particles
32,925 particles
Homogeneous refinement
Homogeneous refinement
Homogeneous refinement
70S
50S
30S
b
FSC
FSC
FSC
Supplementary Figure 9 | Data processing summary of ribosome which were prepared on graphene grid. (a) Workflow of the structure determination. (b) The corresponding FSC and particle orentation plot.

### Slide 12
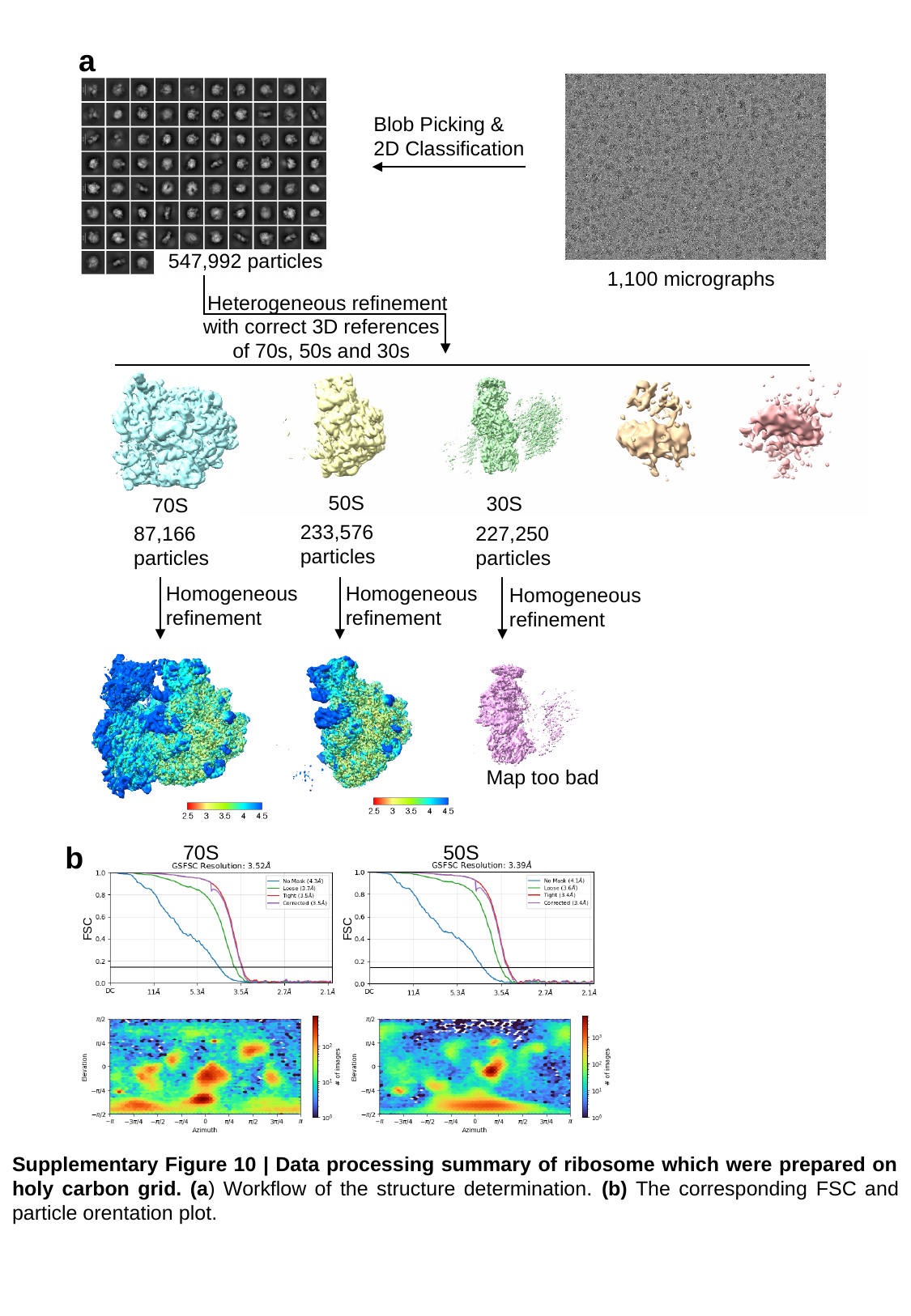

a
Blob Picking &
2D Classification
547,992 particles
1,100 micrographs
Heterogeneous refinement
with correct 3D references of 70s, 50s and 30s
50S
30S
70S
233,576 particles
87,166 particles
227,250 particles
Homogeneous refinement
Homogeneous refinement
Homogeneous refinement
Map too bad
b
70S
50S
FSC
FSC
Supplementary Figure 10 | Data processing summary of ribosome which were prepared on holy carbon grid. (a) Workflow of the structure determination. (b) The corresponding FSC and particle orentation plot.

### Slide 13
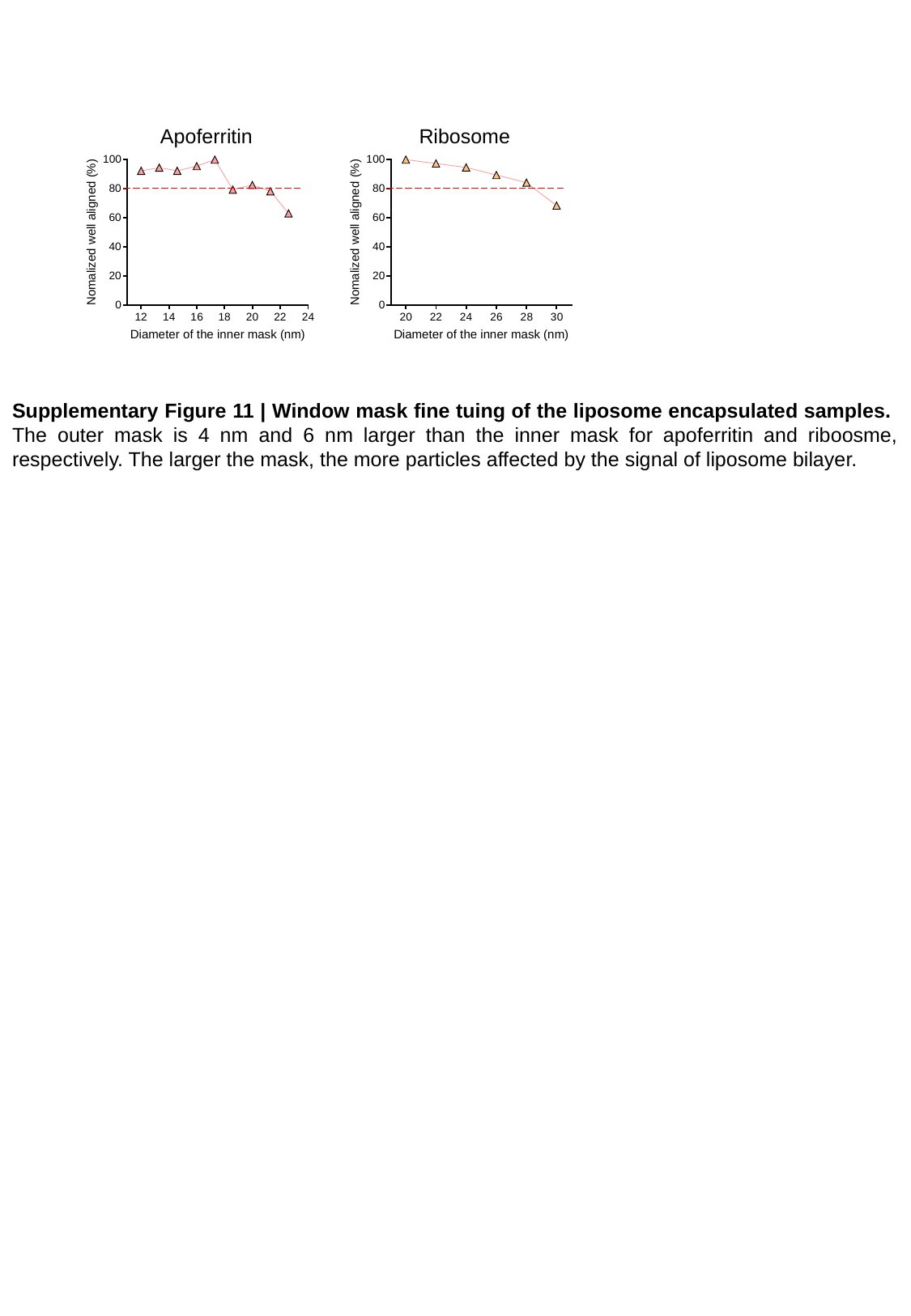

Apoferritin
Ribosome
Supplementary Figure 11 | Window mask fine tuing of the liposome encapsulated samples. The outer mask is 4 nm and 6 nm larger than the inner mask for apoferritin and riboosme, respectively. The larger the mask, the more particles affected by the signal of liposome bilayer.
